# All-optical manipulation of the *Drosophila* olfactory system

**DOI:** 10.1101/2022.02.08.479558

**Authors:** Mirko Zanon, Damiano Zanini, Albrecht Haase

## Abstract

Thanks to its well-known neuroanatomy, limited brain size, complex behaviour, and extensive genetic methods, *Drosophila* has become an indispensable model in neuroscience. A vast number of studies have focused on its olfactory system and the processing of odour information. Optogenetics is one of the recently developed genetic tools that significantly advance this field of research, allowing to replace odour stimuli by direct neuronal activation with light. This becomes a universal all-optical toolkit when spatially selective optogenetic activation is combined with calcium imaging to read out neuronal responses. Initial experiments showed a successful implementation to study the olfactory system in fish and mice, but the olfactory system of Drosophila has been so far precluded from an application. To fill this gap, we present here optogenetic tools to selectively stimulate functional units in the *Drosophila* olfactory system, combined with two-photon calcium imaging to read out the activity patterns elicited by these stimuli at different levels of the brain. This method allows to study the spatial and temporal features of the information flow and reveals the functional connectivity in the olfactory network.

## Introduction

All animals use different sensory modalities to perceive the surrounding environment. This includes vision, hearing, taste, smell, and touch. Between these, olfaction is for many insects the most important sensory modality to ensure survival and reproductive success.

Thanks to recent advances in physiology and genetics, much is known about how *Drosophila melanogaster* encodes odour stimuli in its brain. In fact, it is one of the most used animal models in neuroscience to investigate the logic of olfaction ^1^; the reasons are several: anatomical similarity to the vertebrate olfactory system, reduced brain dimensions facilitating the optical access, precise description of synaptic connections, availability of a vast set of genetic tools for its manipulation, and a large number of paradigms to test olfactory perception.

The majority of olfactory sensilla are located at the distal part of the antenna. Olfactory receptor neurons (ORNs) within the sensilla extend their axons to the first olfactory centres in the brain, the antennal lobes (ALs), the analogue of the olfactory bulbs (OBs) in mammals. Specifically, the ORN axons terminate in spherical substructures, the glomeruli. 43 glomeruli are present in an AL and each one receives input from a single type of olfactory receptors ^2^; local interneurons (LNs) interconnect different glomeruli ^3^ and projection neurons (PNs) forward the glomerular activity to higher brain centres. Odour stimuli produce stereotypical glomerular activation patterns in the ALs, which can be observed via calcium imaging in fruit flies ^4^, but also in other insects like bees ^5^ as well as in the mammalian OBs ^6^. Both spatial and temporal degrees of freedom contribute to odour coding ^7–12^. PN axons connect to higher processing centres like the calyces of the mushroom bodies (MBs) and dorso-lateral regions of the protocerebrum called lateral horns (LHs). This information transfer from ALs to higher centres has been intensively studied ^13–15^. At the MBs level, information is read out by ca. 2000 Kenyon cells (KCs) which are sparsely activated without an evident stereotypical spatial patterning. KCs integrate different combinations of inputs, each from ca. 7 PNs, maximizing stimulus discrimination performance ^16^.

A large effort has currently been made to study connectivity between and within these brain regions. Besides spectacular results showing anatomical connections down to the single synapse ^17^, functional connectivity is studied with various approaches, *e.g.* by combining Ca-imaging with bath-applied ATP pulses ^18^, electrophysiological stimulations and readout of single LNs and PNs ^19^, or by correlating spontaneous activity ^20^. First efforts are made to precisely connect anatomical and functional data^21^. These and other studies gave a good insight into connectivity properties, however, there is one bottleneck in all works which investigate the olfactory system at the network level: how to combine single network node stimulation with whole-brain readout. While in electrophysiological studies, to perfectly shape single neuron stimuli, the response can be recorded only in a few neurons, whole-brain studies use spontaneous activity to determine the resting state connectivity or use odour stimuli which usually elicit responses in a large number of input neurons cross-linked in the AL, typically producing a broad activity pattern.

To shed light on the olfactory network and its coding mechanisms, it would be of great advantage to access also single network nodes and follow the produced activation in space and time. This would help to explore the direct and indirect coupling between single cells, enabling systematic studies on how single odour properties are encoded in this high-dimensional system.

The selective interaction with single neurons became feasible by the experimental implementation of optogenetics, a technique based on the integration of light-sensitive ion channels (opsins) into the cell membrane to modulate its ionic permeability upon illumination with specific wavelengths ^22^. Since the first proof-of-principle experiments ^23–25^, scientists have developed a large toolset of opsins to excite or inhibit single neurons ^26,27^. In combination with spatially and temporally resolved readout, *e.g.* via calcium imaging, this provides a powerful all-optical approach to study neuronal networks ^28,29^.

However, the combined use of optogenetics and calcium imaging is experimentally challenging: besides the costs and complexity of integrating and controlling two laser light sources (one for the activation, one for the readout), the spectral overlap between the molecules responsible for optical activation and calcium activity detection represents one of the biggest problems ^24^. The best performing activity sensors in transgenic animals, the genetically encoded calcium indicators GCaMP6 ^30^, GCaMP7 ^31^, and GCaMP8 ^32^ are overlapping with the most common opsin, channelrhodopsin ChR2 ^24^. Wavelength-shifted opsins and calcium sensors are less efficient ^33^ but still widely chosen to avoid this problem ^34^. Alternative ways to minimize this cross-talk in an all-optical activation and readout system would be the targeting of spatially separated areas for stimulation and imaging or an optimization of laser powers for the two different tasks ^35–39^.

By now, the power of all-optical neuronal manipulation and imaging has been successfully demonstrated in different animal models and various parts of the brain ^40^. The applicability of these setups was extended by advanced stimulus delivery and imaging techniques ^41–45^. Also in Drosophila, the all-optical approach has been applied to various brain regions and neuronal populations ^27,46^. But among those, only a single work concentrated on the olfactory system of adult flies. The experimental approach was, contrary to this work, the stimulation of a single type of ORN which was targeted by selective expression of ChR2 and not by spatially selective illumination ^47^. In a further study in Drosophila larvae, ChR2 was expressed in all ORN, but also here light patterning was not applied, since a response of all ORN was desired ^48^. Thus, to our knowledge, the here presented methods are the first, that show how to dissect the olfactory network in flies via a selective illumination of single glomeruli. However, the benefits of such an approach within the olfactory system have already been demonstrated in fish ^49^ and mice ^50–52^.

To fill this gap and to solve the aforementioned limitations, we implemented optogenetics to selectively stimulate functional units in the *Drosophila* olfactory system via a diode laser and to read out the activation via two-photon fluorescence microscopy providing high resolution and penetration depth. We present new transgenic lines of *Drosophila,* expressing a combination of the best-performing actuators and sensors, the opsin ChR2-XXL ^53^ at the level of the ALs and the calcium indicator GCaMP6 pan-neuronally.

Our results demonstrate that these flies qualify as a reliable and versatile tool to stimulate specific nodes in the primary olfactory network while monitoring the associated neuronal response throughout the brain *in vivo.*

## Results

### 1. Antennal lobe stimulation and olfactory pathway connectivity

To validate our all-optical system and transgenic model, we monitored the entire fly brain activity, stimulating only one antennal lobe.

The correlations between the response signals of different brain areas were calculated (*Fig. 1b*), including the left and right ALs, the left and right MBs and the left and right posterior brain (superior medial protocerebrum) (*Fig. 1a*). In all cases, correlations are higher for the left side stimulation, which is likely influenced by the specific area that was targeted. Looking at the ipsilateral correlations, the largest ones are those between the stimulated AL and the ipsilateral MB (*r* = 0.79±0.09 (mean±sem) for the left, *r* = 0.51±0.10 for the right side stimulation); they then reduce between stimulated AL and posterior brain (*r* = 0.60±0.32 for the left and *r* = 0.17±0.06 for the right side). Of major interest are the contralateral couplings which are relatively low between antennal lobes (*r* = 0.30±0.30 under left stimulation, *r* = 0.13±0.29 under right stimulation) and increase substantially between ALs and contralateral MBs (*r* = 0.62±0.20 under left stimulation, *r* = 0.38±0.24 under right stimulation) and between both MBs (*r* = 0.60±0.25 under left stimulation, *r* = 0.61±0.09 under right stimulation).

**Figure 1.**
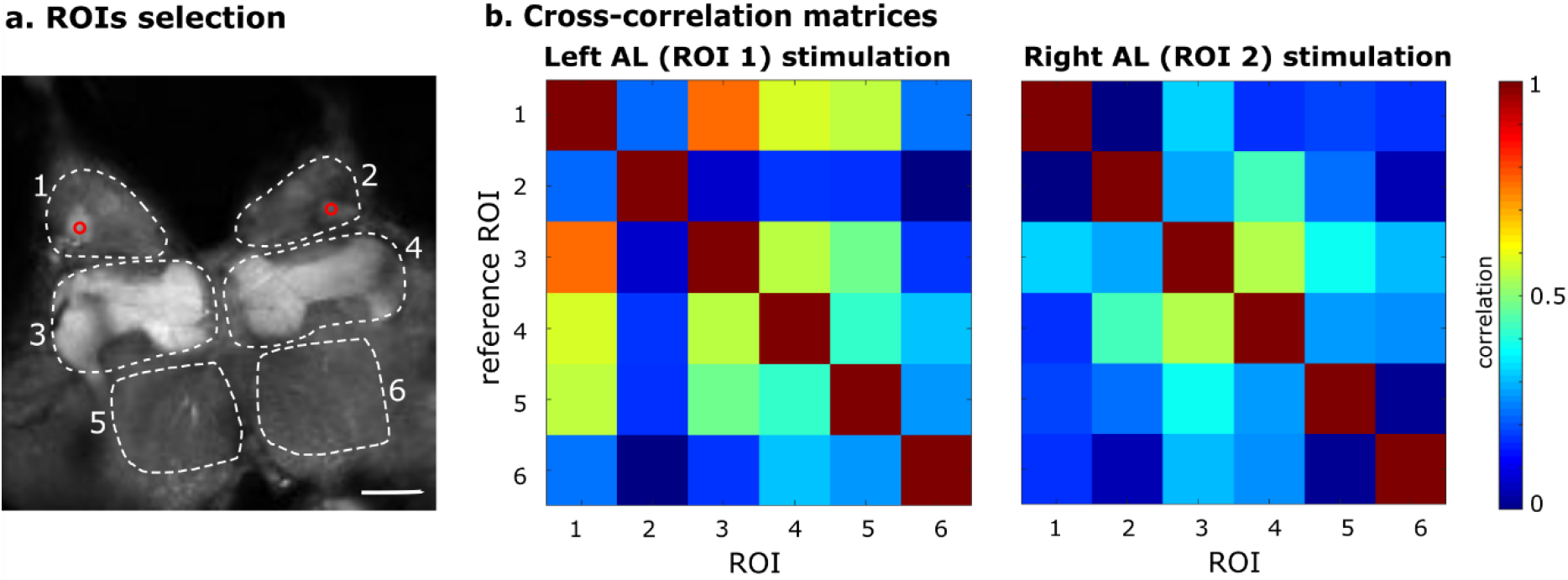
Whole brain imaging. a) Cross-section fluorescence image of a whole brain, with different ROIs delimiting brain areas involved in olfactory processing (1,2 ALs; 3,4 MBs; 5,6 posterior brain areas). The red circles represent the two different stimulation points (in two different experiments). Scale bar 80 μm. b) Cross-correlation analysis between different areas of the fly brain, following an antennal lobe stimulation. The Pearson’s correlation coefficient is represented as a colour code. The results show an average over 3 stimulus repetitions.

**Figure 2.**
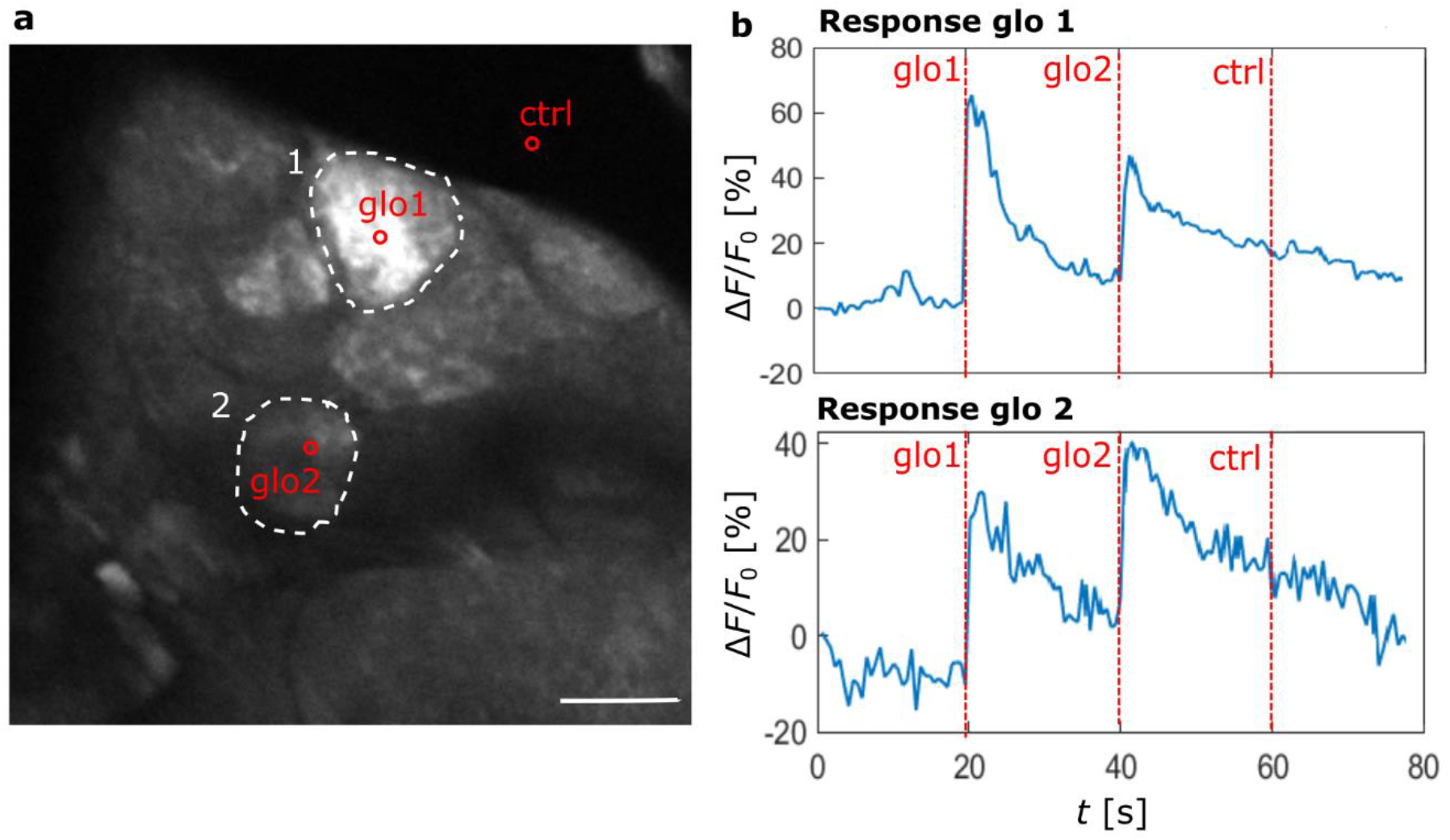
Glomeruli stimulation. a) Cross-section image of an AL, highlighting two glomeruli (white ROIs) and three stimulation points (red circles). Scale bar 20 μm. b) Temporal response curves averaged over the two different ROIs (glomerulus 1 above and glomerulus 2 below); red vertical lines indicates the times of stimulation: the first stimulation targets glomerulus 1, the second glomerulus 2, and the third a point outside the AL (ctrl). The interval between stimuli is 20 s.

### 2 Single glomerulus stimulation and power threshold

To explore the resolution of our setup and test the feasibility to solely stimulate neurons within a single glomerulus of interest, we stimulated three spots consecutively, targeting two different glomeruli and, as a control, an area outside the AL.

The temporal response curves manifest that the stimulus elicits a strong response signal in the targeted glomerulus while stimulating the control region outside the AL does not elicit any glomerular activity. However, when targeting one glomerulus, a reduced response was visible also in the other, non-targeted glomerulus. The comparable distance to the control suggests that this might not be a direct activation due to a limited resolution of the activation pattern, but rather an excitatory coupling between glomeruli via local neurons ^54^.

To clarify this, we tested neuronal responses in the neighbouring regions of the stimulated glomerulus. A single glomerulus was repeatedly activated with increasing blue laser power. The time-dependent response curve shows a strong activation of this glomerulus already for the lowest power *p*_1_ = 0.02 mW (orange curve in *Fig. 3b*). An increase of the activation by exposure to higher powers seems to lead always to a similar jump of the signal; the total increase derives from the fact that previous stimulus activity persists. After *p*_5_ a saturation effect can be observed. The nearest neighbour glomerulus (blue curve in *Fig. 3b*) shows no activation for the lowest power *p*_1_, followed by a slow increase with increasing stimulus intensity until *p*_4_ when its response becomes comparable to the targeted glomerulus. An even clearer picture is drawn by a seeded cross-correlation map (*Fig. 3c*) that shows an extremely coherent signal within the stimulate glomerulus, with *r* ≈ 1 that abruptly drops to *r* ≈ 0 at the glomerular boundary, proving the activation is limited to a single glomerulus for stimulation with *p*_1_.

**Figure 3.**
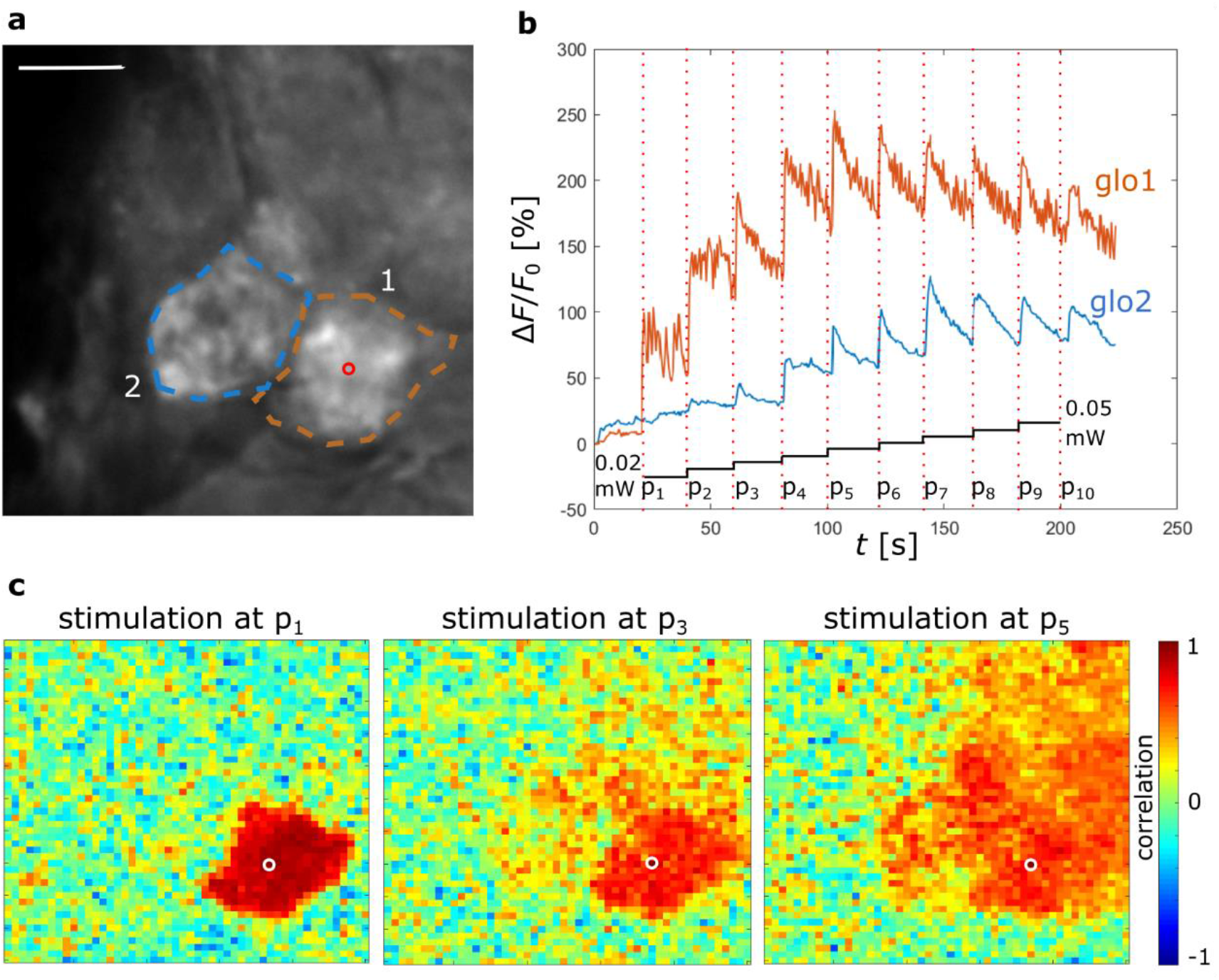
Increasing power stimulation. a) ROIs selection: glomerulus 1, orange ROI; glomerulus 2, blue ROI; the stimulation point in glomerulus 1 is marked by a red circle, scale bar 20 μm. b) Temporal response profile averaged over each of the two ROIs during 10 stimulations with ascending blue laser power from *p*_1_ = 0.02 mW to *p*_10_ = 0.05 mW; black steps show the progressive increase in power for the different 10 stimulations. c) Three examples of seed-based correlation maps with respect to the stimulation point for three stimulations powers *p*_1_, *p*_3_, and *p*_5_.

When the activation power is increased, a progressive extension of activation to neighbouring glomeruli can be observed (*Fig. 3c*).

### 3. Elicited coupled oscillations

Our initial findings also showed interesting phenomena that go beyond the activation of a tonic response limited to the stimulus period which then propagates to higher brain centres.

The example reports the activation of a single glomerulus (glomerulus 1 in *Fig. 4a*) which shows spontaneous activity before the stimulus (*Fig. 4d,* blue line). After the stimulation, this glomerulus maintains its oscillatory activity, although with slightly increased intensity. However, most interestingly, after the stimulation, other glomeruli start to respond in high synchrony to the targeted glomerulus (see correlation matrices of *Fig. 4b*), partially with very high amplitudes that keep increasing until long after the stimulus. The synchrony of these oscillations is nicely visible in *Supplementary Video S1.*

**Figure 4.**
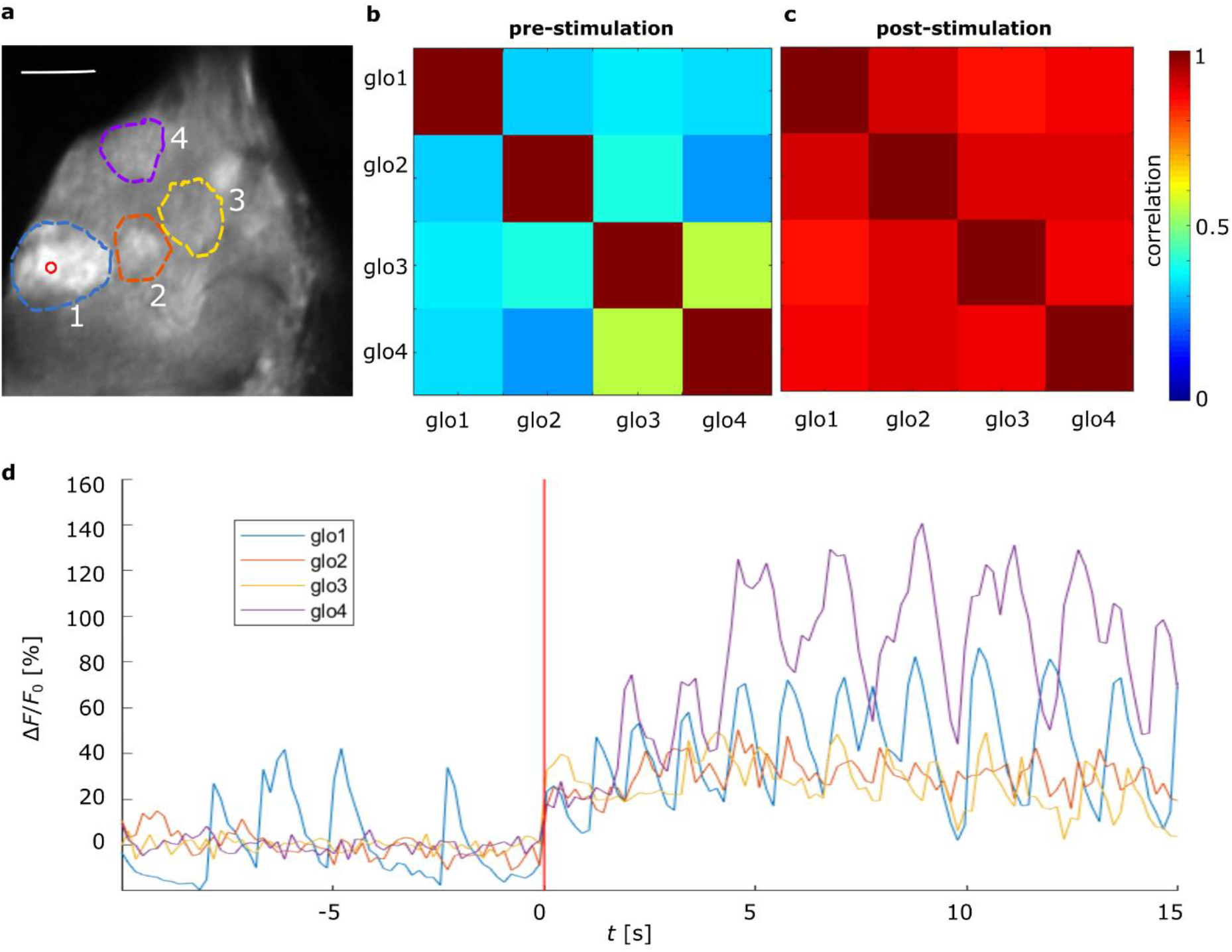
Oscillatory glomerulus stimulation. a) Antennal lobe with ROIs on different glomeruli, and an activation point within glomerulus 1 labelled by a red circle, scale bar 20μm. Cross-correlation matrices in time windows of 10 s before (b) and after (c) the stimulation. d) Temporal activity profiles before and after the 200 ms stimulation, the latter is marked by the red vertical line.

To investigate the dynamics of these oscillations more in detail, a time-frequency analysis of coherence was performed (*Fig. 5*). The example shows the coherence between the spontaneously oscillating glomerulus that was stimulated (glomerulus 1 in *Fig. 4a*) and its neighbouring glomerulus (ROI 2 in *Fig. 4a*).

**Figure 5.**
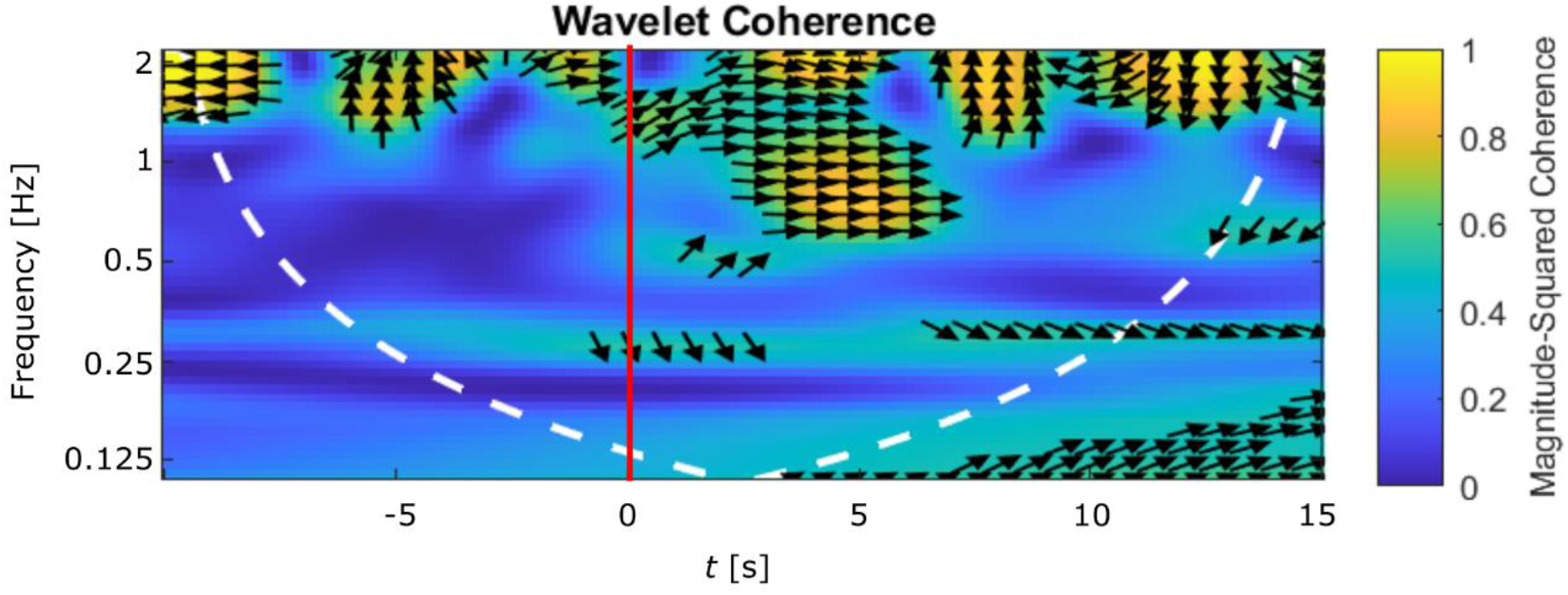
Time-frequency analysis of Exp. 3. Magnitude-squared wavelet coherence between the targeted glomerulus 1 and glomerulus 2 of Fig. 4a. Stimulus at the red vertical line; the white dotted line represents the cone of influence and the arrow directions indicate the phase shift between both signals.

The results suggest fluctuating coherences above 1 Hz with an arbitrarily fluctuating phase before the stimulus. After the stimulus, a new component comes up between 0.5 and 1 Hz where both signals are well in phase, manifested by the right-pointing vector, suggesting coherent oscillations around this frequency. This is the frequency band in which the collective oscillations are observed (*Fig. 4d, Supplementary Video S1).*

## Discussion

In this work, we propose and implement an all-optical approach to study the *drosophila* olfactory system. We provide all the necessary tools to dissect this neuronal network, with applicability on different scales, from the entire brain to single network nodes.

To optimize neuronal activation and response detection efficiency, two of the best performing molecules (ChR2-XXL and GCamp6) were used, which so far were rarely expressed in combination, given their spectral overlap.

The method was first tested at the scale of the entire brain while stimulating a small region within one single antennal lobe. The elicited response patterns were analyzed by looking at the temporal correlations between the activity in different brain regions. The correlation amplitudes (*Fig. 1b*) confirm the well-known pathways of the olfactory information, *i.e.* information is directly forwarded by projection neurons into the ipsilateral mushroom body and lateral horn. This reflects in correlations of 0.8/0.5 (left/right stimulation, respectively) between AL and MB of the same hemisphere. An important difference is visible in the homotopic connectivity: some correlations between contralateral antennal lobes are present, but with smaller amplitudes of 0.3/0.1 (left/right stimulation, respectively). This confirms how in *drosophila* ^55,56^, differently from other insect species ^57^, ORNs target also the homotopic glomerulus in the contralateral AL. The fact that in these regions the correlations are not as high as the ipsilateral ones with the MBs is likely due to the different local network structures created by modulatory interneurons that couple ipsilateral glomeruli. In honey bees, it was shown that also the antennal lobe output signals are therefore not fully bilaterally symmetric ^58^. Contralateral correlations increase substantially between MBs to 0.6/0.6 (left/right stimulation, respectively), confirming the enhanced bilateral coupling of these neuropils ^59^.

To reveal more details on these couplings, the system needs to work at an increased spatial and temporal resolution, which we tested thereafter. Experiments on targeting individual glomeruli demonstrated that the choice of laser power is the most crucial factor for the selectivity of glomerular stimulation (*Fig. 3*). With high spatial resolution and precise control of the activation laser power, a correlation measurement shows how to find an optimal threshold to elicit activation perfectly confined to the targeted glomerulus (*Fig. 3c*). At the same time, it showed how the massive coupling within each glomerulus produced perfect correlations across the glomerular area. The abrupt decay to zero at the glomerular boundaries is proving that the above mentioned spectral overlap between opsin and the calcium-sensor could be well controlled by keeping the two-photon laser intensity reasonably low. The unwanted re-excitation of the opsin during the imaging phase would have been visible as a background in the correlation analysis.

When the activation laser power was successively increased, the neuronal activation reached over into neighbouring glomeruli, likely due to the laser intensity surpassing the activation threshold in more and more neighbouring areas. Although this spillover has to be avoided in protocols targeting single glomeruli, there are also possible applications for it. It has been shown that glomeruli responsive to specific odours tend to cluster topographically; moreover, similar chemical classes of odours and even response profiles of odours with equal hedonic valence highly overlap^60^. It follows that, just by intensity tuning, a single optogenetic activation could produce activity patterns mimicking specific odours, broader odour classes, or even odour valence properties. This may also allow comparing the fundamentally different odour coding strategies of combinatorial coding versus single glomerulus labelled lines ^15,61^ by directly transitioning between these patterns. The corresponding laser powers depend on various parameters, among those the sample staining, the imaging depth, the type and the position of the targeted glomeruli: thus, a calibration needs to be performed before each experiment by the presented experimental scheme.

Besides measuring the static coupling between neural network nodes based on average response amplitudes, we showed that this setup allows accessing also dynamical features of this coupling. The presented example (*Fig. 4* and *Supplementary Video S1*) showed how optical stimulation hardly changed the activity in a stimulated glomerulus that was already showing a spontaneous response, but at the same time caused a long-lasting coupling between the glomerulus and its neighbours (which again might have been due to coupling via excitatory interneurons ^54^). This highlights the potential of this approach for studying also the dynamical coupling under well-defined conditions, limiting the excitation to a single glomerulus. An odour stimulus would have likely activated several glomeruli, making it impossible to identify the origin of such synchronization.

These are proof-of-principle experiments that show how optogenetics can be used as a powerful tool to study signal propagation and functional connectivity in the *Drosophila* olfactory system. It will be possible to reconstruct the functional connectome of different brain regions with odour-independent optogenetic stimulation of single network nodes. It allows, moreover, to follow the dynamics of information propagation from single, well-defined sources.

## Methods

### Drosophila lines

All fly lines used in this study were reared on standard cornmeal-agar medium with yeast, at 20°C in 60% humidity-controlled chambers under 12 hours light/dark cycles. The fly lines were obtained from the Bloomington Drosophila Stock Center: *orco-GAL4* (#26818); *UAS-ChR2-XXL* (#58374); *nSyb-LexA* (#52247); *LexAop-GCaMP6m* (#44275).

The flies used in the experiments were obtained by combining both Gal4-*UAS* and LexA-*LexAOP* binary systems into one fly, simultaneously performing two manipulations of gene expression *in vivo.* In our tests we used 5 days old flies *nSyb-LexA/UAS-ChR2-XXL; LexAop-GCaMP6m/orco-GAL4* which express the channelrhodopsin in olfactory sensory neurons and the calcium sensor in all the neurons of the brain.

Correct and precise expression of transgenes was checked by performing immunohistochemistry on dissected brains, following a standard protocol ^62^ using primary antibodies specific to GCaMP (chicken anti-GFP, Thermo Fisher Scientific) and ChR (mouse anti-ChR2 supernatant 1:1, 15E2, mfd Diagnostics). Secondary antibodies were goat anti-chicken Alexa Fluor-488 and goat anti-mouse 1:250, respectively (Thermo Fisher Scientific). *Fig. 6b,c,d* show the expression levels in the fly brain. The opsin expression is limited to the antennal lobes (*Fig. 6b,* magenta in *Fig. 6d*), the calcium sensor is found in all neurons (*Fig. 6c,* green in *Fig. 6d*).

**Figure 6.**
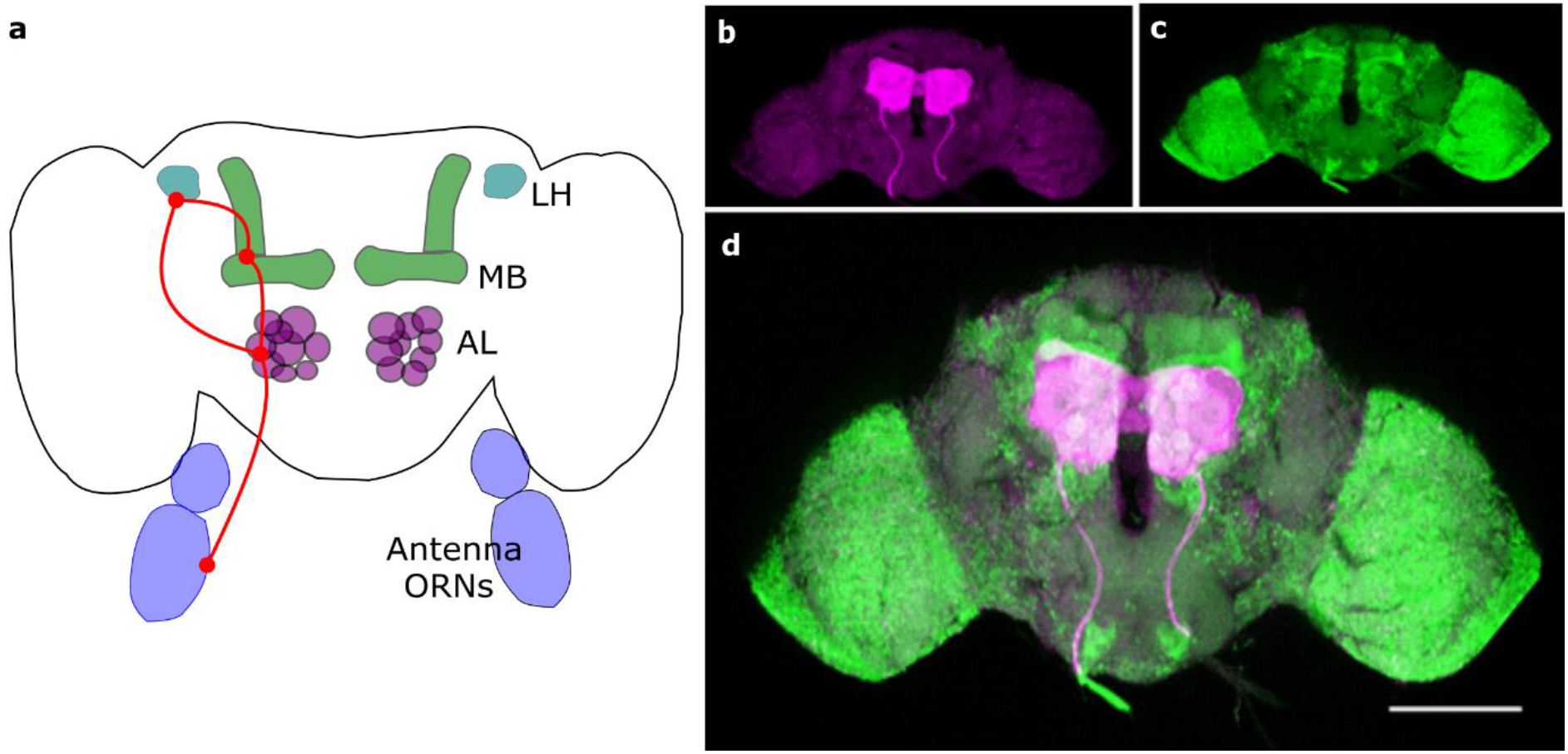
Fly brain and molecules expression. a) Scheme of the fly olfactory system. Odours are received by the odour receptor neurons (ORNs) at the level of the antennae. The activation is passed to the antennal lobes (ALs) and further to the mushroom bodies (MBs) and the lateral horns (LHs). (b) Images of a dissected and fixed fly brain, marked with antibody anti-ChETA (b, magenta) for ChR2-XXL identification and anti-GFP (c, green) for calcium sensor GCaMP identification. The opsin is expressed in the antennal lobes, GCaMP pan-neuronally. c) Merge of (b) and (c). Scale bar: 100 μm.

### Drosophila preparation

Sample preparation and drosophila brain exposure are adapted from a well-established procedure ^63,64^.

1. Custom-made plexiglass mounting blocks (*Fig. 7a*) are used to allow easy mounting and dissection of the fly, blocking the animal by the neck. A copper disc (125 μm slot, Copper 3.05mm, Agar Scientific) is glued and levelled on the block to create a “collar”, with the slit centred above the hole of the block. The flaps on both sides of the slit are folded down to stay close to the block surface.
2. Flies of the appropriate genotype are anaesthetized by cooling on ice, females are chosen for their bigger dimensions. Holding the fly by the wings, it is introduced into the mounting block by the neck and rotated until it is looking downwards. When the head is levelled with the top of the block, the fly is blocked with a cactus spine above the proboscis and a drop of glue (Super Attak, Loctite) on the sides of the head (*Fig. 7c*).
3. Fly antennae are pushed away from the head with a thin wire (attached to a plastic U-shaped coverslip, across the top of the “U”) (*Fig. 7b* and *7d*). This wire is inserted in the cuticular fold between head and antennae and the screws built into the mounting block (*Fig. 7a*) are used to gently adjust the wire position.
4. An antennal shield composed of a circular slot in a not-sticky piece of tape surrounded by a plastic coverslip is placed on the top of the head. The slit has the same dimensions as the head, not extending beyond the eyes. This shield prevents the preparation from leaking, keeping the antennae completely dry (*Fig. 7e*).
5. The gap between the tape and the cuticle is sealed with two-component silicon (Kwik-Sil, WPI), in a thin layer; a drop of *Drosophila* Ringer’s solution (130 mM NaCl, 5 mM KCl, 2 mM MgCl2, 2 mM CaCl2, 36 mM saccharose, 5 mM HEPES, pH 7.3) is then placed on the head.
6. The head cuticle is cut using a sapphire blade (Single/Double Edge Lancet, 1.00 mm wide, angled 45°, WPI), carving along the borders of the eyes and across the ocelli on the back. After removing gently the piece of cuticle, glands and trachea are removed with fine forceps (Dumont Tweezers #5SF, 0.025 × 0.005mm, WPI), and the fly’s brain is ready to be imaged.

**Figure 7.**
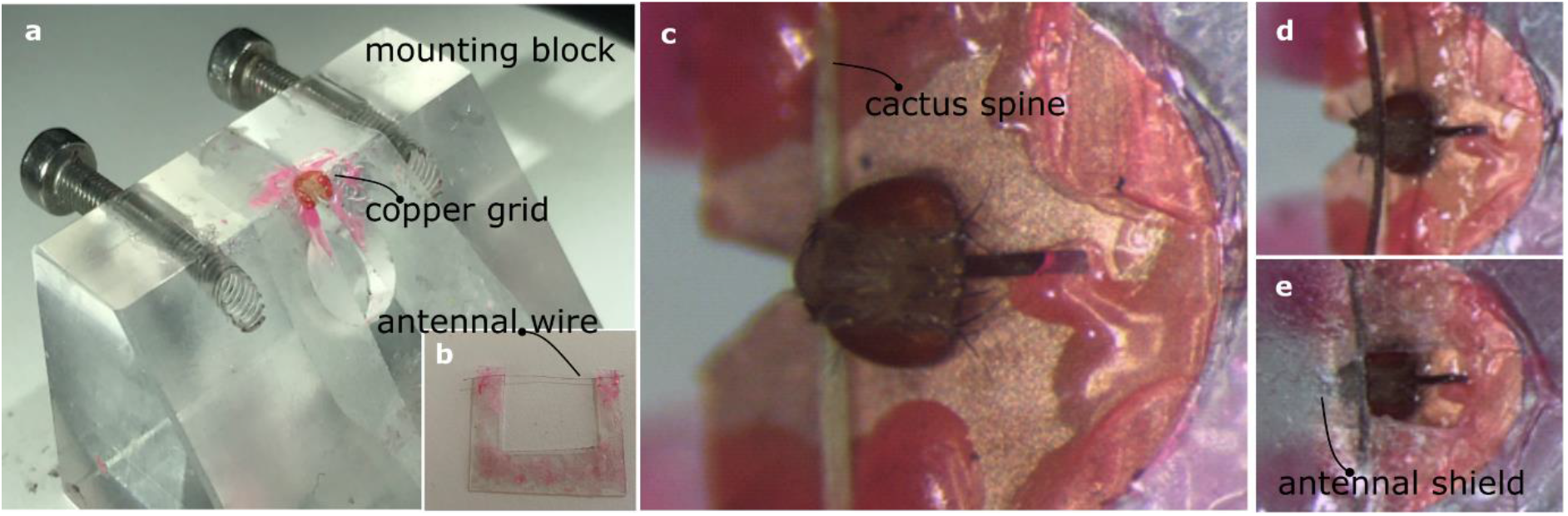
Drosophila preparation. a) Picture of the plexiglass mounting block, with the copper disc on the top. (b) Mounted wire that is used to position the antennae. c) The drosophila is blocked with a cactus spine after it’s inserted in the copper disc by the neck. d) The antennae are pushed by the wire (b). e) antennas are covered with a plastic shield.

### Microscopy setup

Our optical setup consists of a two-photon microscope (Ultima IV, Bruker) combined with a femtosecond pulsed Ti:Sa laser (MaiTai DeepSee, Spectra-Physics), tunable between 690-1040 nm. The output power is controlled by a Pockels cell. Two-photon imaging of GCaMP worked best at a wavelength of 940 nm.

The beam is scanned in the plane by galvo-mirrors and in height by a fast piezo scanner (150 μm travel range) and a stepper motor (travel range of 25 mm), which move the water-immersion objective (20× NA 1.0, Olympus). Fluorescence is separated by a dichroic beam-splitter (660 nm, ChromaTechnology), filtered by an IR-blocking filter (< 650 nm) and divided by dichroic beam-splitter (575 nm) into two channels equipped with two different band-pass filters (607/45 nm and 525/70 nm, ChromaTechnology) and detected by multi-alkali PMT detectors (Hamamatsu). The complete setup is placed on an air-damped optical table (RS4000 Newport).

To stimulate the opsin, an additional diode laser emitting at 473 nm with a maximum output power of 100 mW (iBeam Smart 473 nm, Toptica Photonics) is overlapped with the imaging beam. In the experiment, the output power of the blue laser is always less than 0.05 mW and the stimulations are point-like, with a duration of 200 ms. All reported power values are measured over the entire beam after the objective. A notch filter (405/473-488/NIR m) blocks the blue light in the fluorescence detection arm. The blue laser spot dimensions (*i.e.* the limit for single point stimulation) are estimated by bleaching a homogeneous fluorescent sample (fluorescent marker pen on coverslip). The obtained point spread function doesn’t exceed 5×5 μm^2^ in the focal plane and has an extension of less than 20 μm along the axial direction. The activation scanning and the imaging beam is handled by the same galvo mirror pair, switching within 1 ms between the continuous 2D line-by-line scanning during imaging and random access scanning of a series of predefined activation points. Moreover, an optional spiralling over these points can be set to increase the exposed areas. As a compromise between spatial and temporal resolution, the imaging experiments are performed using 128×128 pixels with a 5 Hz repetition rate. The microscope is controlled by the software PrairieView (Bruker).

### *In vivo* imaging

#### 1. Antennal lobe stimulation and olfactory pathway connectivity

We monitored the entire fly brain activity, in response to the stimulation of a single antennal lobe; activation was performed either within the left AL or within the right one. The stimulation was a single-point blue-laser activation (red circle in *Fig. 1a*), with a power < 0.03 mW for a duration of 200 ms. Activity propagation was quantified via correlation analyses between different brain areas (see below for a detailed description of the analysis). All results were averaged over three stimulus repetitions.

#### 2. Single glomerulus stimulation and power threshold

In a first experiment, three spots were stimulated consecutively, targeting two individual glomeruli and an area outside the AL. The sequence of stimulation was the following: glomerulus 1, 2, and a control point outside the AL, with a delay of 20 s between stimuli (*Fig. 2*). The stimulations were point-like, with a power *p* < 0.03 mW for a duration of 200 ms.

In a second experiment, a single glomerulus was repeatedly activated with increasing blue laser power (corresponding to powers ranging from *p*_1_ = 0.02 mW to *p*_10_ = 0.05 mW; *Fig. 3*). The response intensity was measured with high resolution in the surrounding of the glomerulus. Seeded cross-correlation maps were constructed to monitor the correlations of different AL areas with the activation point, at increasing power (see the following paragraph for detailed analysis description).

#### 3. Elicited coupled oscillations

We activated a single glomerulus (glomerulus 1 in *Fig. 4a*) that was showing a spontaneous oscillatory activity before the stimulus. The stimulation was point-like, with a power < 0.03 mW for a duration of 200 ms. A time-frequency analysis of the magnitude-squared coherence between the oscillatory signal of the targeted glomerulus and the surrounding ones was performed (*Fig. 5*).

### Analysis of fluorescence activity and data correlation

Neuronal responses to stimulation are measured as the change in fluorescence Δ*F* with respect to a baseline value *F*_0_ (average fluorescence before stimulation). All response data are normalized with respect to this baseline to allow comparison between different samples and subjects. The formula used for this purpose is: Δ*F*/*F*_0_ [%] = (*F - F*_0_)/*F*_0_*100.

Image processing and data analysis were performed using custom scripts in Matlab (R2019, MathWorks). These include, besides the above-mentioned post-processing, correlation analyses between time series (*Fig. 1b, 4b*) and seed-based spatial correlation analyses between an activation point and its surrounding (*Fig. 3c*). Correlation analyses are based on *corrcoef* Matlab functions. Furthermore, a wavelet coherence analysis was performed (*Fig. 5*) to obtain a time-frequency plot of the magnitude-squared coherence between two signals (colour-coded) plus the time-dependent phase between the two signals (coded by arrows, horizontal forward-pointing meaning completely in phase). This analysis is based on *wcoherence* Matlab function.

All the afromentioned analysis were implemented into a custom graphic user interface (*Supplementary Fig. S1, S2* and *S3*).

## Additional information

### Availability of materials and code

The program code and data are freely available at https://github.com/MirkoZanon/OptoFluorecenceAnalysis

### Competing interests

The authors declare no competing interests.

### Authors’ contributions

M.Z. performed the experiments and the analysis. D.Z. engineered the Drosophila lines and performed immunohistochemistry. A.H. supervised the project. All the authors contributed to the design of the experiments and to the final version of the manuscript.

## Notes

### Competing Interest Statement

The authors have declared no competing interest.

