## Supplementary Material for "All-optical manipulation of the *Drosophila* olfactory system"

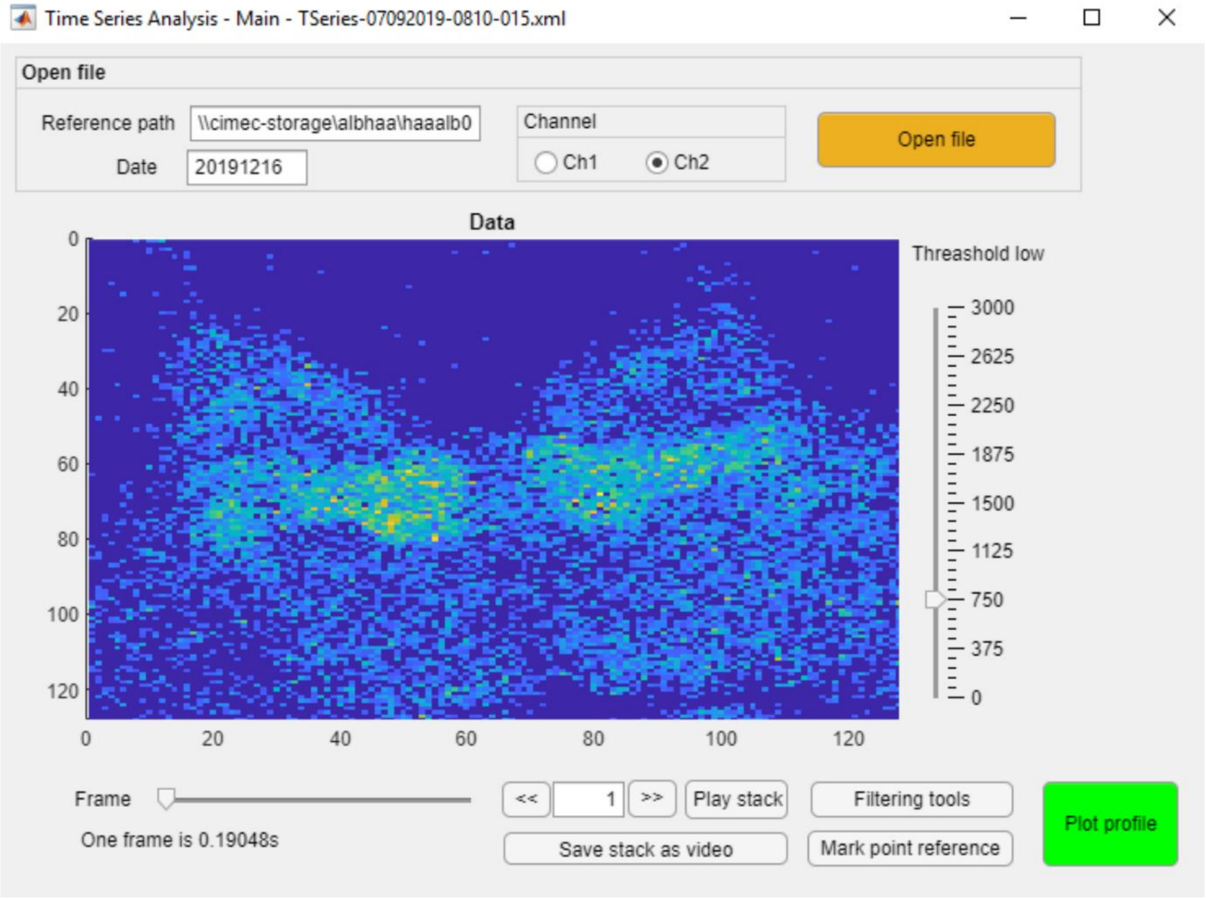


**Supplementary Figure S1** *Analysis GUI – Loading tool*. Example of the custom Matlab graphic user interface used to elaborate and analyse the fluorescence data following optogenetic stimulation. This window allows to load the time series, perform noise thresholding and access all the other tools of the custom software.

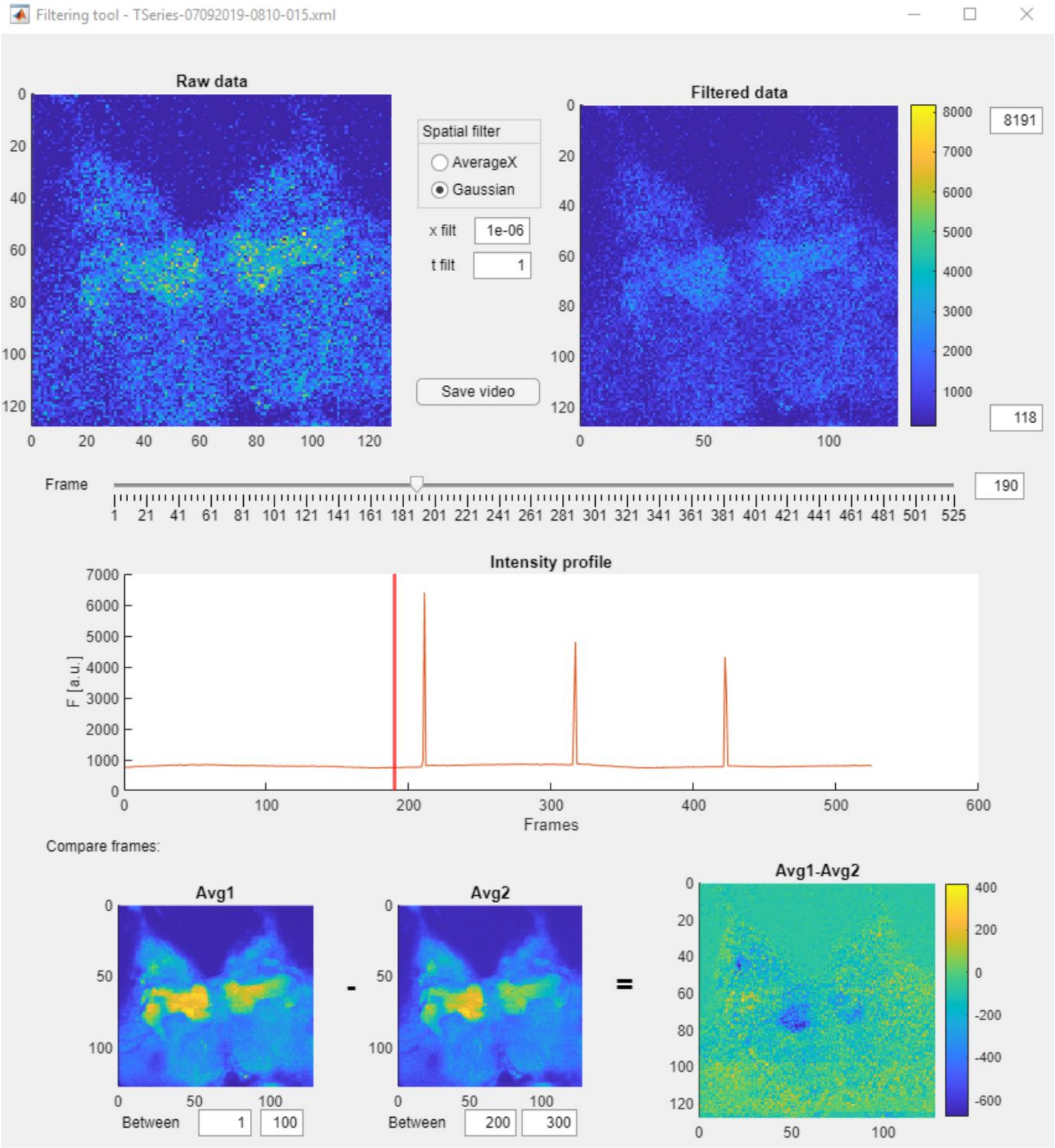


**Supplementary Figure S2** *Analysis GUI – Filtering tool*. Example of the custom Matlab graphic user interface used to elaborate and analyse the fluorescence data following optogenetic stimulation. This window allows to perform spatial and temporal filtering of the time series and compare averages of different time regions.

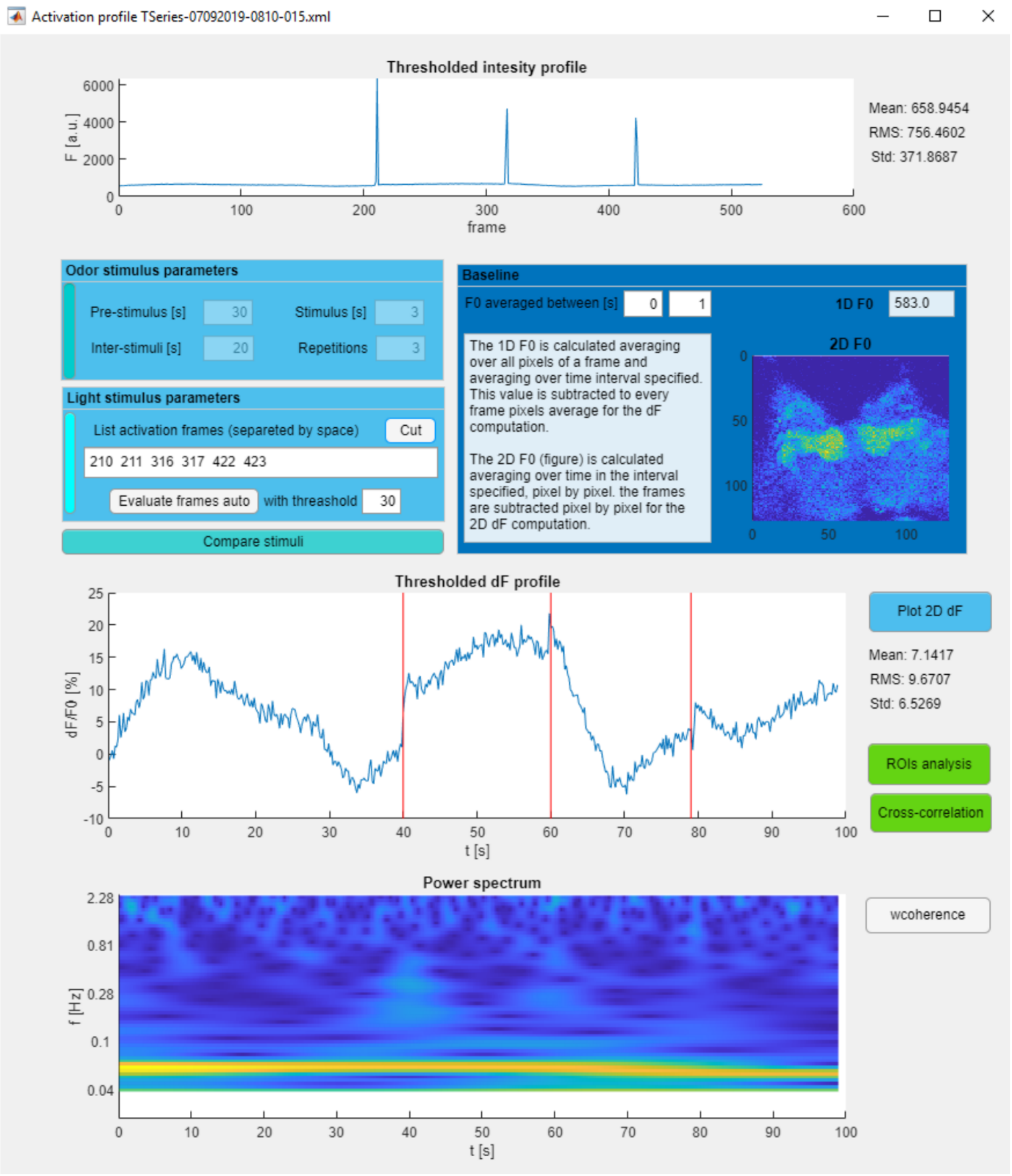


**Supplementary Figure S3** *Analysis GUI – Intensity and correlation tool*. Example of the custom Matlab graphic user interface used to elaborate and analyse the fluorescence data following optogenetic stimulation. This window allows to reconstruct the fluorescence intensity profile, eliminate optogenetics blue light saturations during activation and perform time-frequency analysis. It also guides the user through all the passages to calculate correlations and perform other analysis between custom ROIs.

**Supplementary Video S1** *Real-time recording from Experiment 3*. Two-photon fluorescence signal of the neuronal activity of an antennal lobe during an optogenetic stimulation experiment (*Experiment 3*). The targeted glomerulus shows an oscillatory spontaneous activity even before the blue light stimulation (visible in the video from the photomultiplier saturation). After optogenetic stimulation, an in-phase oscillatory response is induced also in the neighbouring glomeruli.
